# Clathrin-mediated endocytosis-independent function of VAN3 ARF GTPase Activating Protein at the plasma membrane

**DOI:** 10.1101/2021.11.17.468981

**Authors:** Maciek Adamowski, Jiří Friml

**Author notes:** Author contributions: M.A. and J.F. designed research, analyzed data and wrote the manuscript. M.A. performed research.

## Abstract

ARF small GTPases in plants serve important cellular functions in subcellular trafficking and developmental functions in auxin-mediated patterning of the plant body. The *Arabidopsis thaliana* ARF regulator ARF-GAP VAN3 has been implicated to act at the plasma membrane (PM) and linked functionally to the clathrin- and dynamin-mediated endocytosis. Here we re-evaluated the localization of VAN3 at the PM and its function in endocytosis. Using Total Internal Reflection Fluorescence microscopy we observed remarkably transient associations of VAN3 to the PM at discrete foci, however, devoid of clathrin, the dynamin isoform DRP1A, or the ARF regulator GNOM, which is also involved in a developmental patterning function mediated from the PM. Clathrin-coated pits are abundant and endocytosis appears to proceed normally in *van3-1* knockout mutant. In turn, post-translational silencing of clathrin expression indicates that the localization of VAN3 at the PM depends on clathrin function, presumably on clathrin-mediated endocytosis.

## Introduction

ARF small GTPases function in the endomembrane system of the cell as recruitment platforms for various effector proteins, including vesicular coat components (reviewed in Donaldson and Jackson, 2011; Jackson and Bouvet, 2014; Yorimitsu et al., 2014; Singh and Jürgens, 2017). ARFs are activated, dock into membranes, and recruit their effectors transiently and reversibly due to their ability to switch between GTP- and GDP-bound forms, which corresponds with conformational changes in the protein. The ARF switches are controlled by two classes of ARF regulators, ARF-GEFs (ARF Guanine nucleotide Exchange Factors) and ARF-GAPs (ARF GTPase Activating Proteins). In plants, studies indicate functions of the ARF machinery in secretory and endocytic activity of the cell (Teh and Moore, 2007; Richter et al., 2014, 2007; Naramoto et al., 2010; Kitakura et al., 2017; Tanaka et al., 2009, 2013, 2014; Xue et al., 2019), while on the organismal level, some of the components play prominent roles in developmental patterning mediated by the plant hormone auxin (Mayer et al., 1993; Steinmann, 1999; Richter et al., 2010; Sieburth et al., 2006; Koizumi et al., 2005; Adamowski et al., 2021a).

Fifteen proteins with predicted ARF-GAP domains exist in *Arabidopsis thaliana*, designated AGD1-15 (for Arabidopsis GAP Domain 1-15; Vernoud et al., 2003; Singh and Jürgens, 2017). These fall into several subclasses based on the presence of other domains in each protein. The evolutionarily conserved ACAP (Arf GAP with coiled coil, ANK repeat and PH domains) class of ARF-GAPs is characterized, beside an ARF-GAP domain, by the presence of an N-terminal coiled coil BAR (Bin, amphiphysin and Rvs) domain, sensitive to membrane curvature, followed by a PH (Plecstrin homology) domain, which binds specific membrane lipids, and at the C-terminus by Ankyrin repeats, often involved in protein-protein interactions. This class is represented in *A. thaliana* by AGD1-4, of which AGD1-3 conform to this domain composition exactly, while AGD4 lacks the N-terminal BAR domain. AGD1-4 ARF-GAPs have been functionally characterized based on the isolation of *vascular network defective3* (*van3*), *scarface (sfc)*, and *forked2 (fkd2)* mutants, deficient in *AGD3*, in forward genetic screens aimed at the identification of mutants with aberrant patterns of leaf vascular networks (Koizumi et al., 2005, 2000; Deyholos et al., 2000; Sieburth et al., 2006; Steynen and Schultz, 2003). *agd3* mutants exhibit fragmentation of the vascular pattern in cotyledons and leaves, concomitant with a fragmentation of expression domains of PIN-FORMED1 auxin efflux carrier (PIN1; reviewed in Adamowski and Friml, 2015) earlier during vascular network development (Scarpella et al., 2006). Higher order mutants in *agd1-4* group exhibit an increase in the severity of vascular patterning defects of *agd3* (Sieburth et al., 2006). The developmental roles of AGD3/SFC/FKD2/VAN3 and its homologues extends beyond vascular network development in cotyledons and leaves, as evidenced for instance by the observation that *agd3* mutants do not develop into the generative stage (Deyholos et al., 2000), or that higher order mutants exhibit patterning defects in lateral root formation and cotyledon development (Naramoto et al., 2010).

Several lines of evidence support the notion that on the molecular level, the function of AGD3/SFC/FKD2/VAN3 (VAN3 in the following) is associated with clathrin-mediated endocytosis (CME). First, as discussed, VAN3 contains a BAR domain, which likely gives the protein curvature-sensing abilities, and as such, is expected to aid its targeting to high-curvature membranes such as those present at forming vesicles (Simunovic et al., 2015; Renard et al., 2018; Frost et al., 2009). Second, fluorescent protein fusions of VAN3, whose expression complements the mutant phenotype, localize to the plasma membrane (PM), both in seedling root apical meristems (RAMs) and in more functionally relevant tissues of the cotyledon (Naramoto and Kyozuka, 2018). Regulators of VAN3 function, VAB/FKD1 (VAN3-BINDING PROTEIN/FORKED1), CVP2 (COTYLEDON VASCULAR PATTERN2), as well as its homologue CVL2 (COTYLEDON VASCULAR PATTERN2-LIKE2) localize to the PM as well (Naramoto and Kyozuka, 2018; Naramoto et al., 2009; Hou et al., 2010; Carland et al., 1999; Carland and Nelson, 2009). Third, in yeast two hybrid screening, VAN3 was found to physically interact with DRP1A (DYNAMIN RELATED PROTEIN 1A; Sawa et al., 2005), a plant-specific isoform of dynamin, a mechanoprotein involved in the separation of formed coated vesicles from source membranes (Collings et al., 2008; Antonny et al., 2016). Finally, fluorescent live imaging-based studies report an association of VAN3 with clathrin-coated pits (CCPs) at the PM, and a reduction of the uptake of the fluorescent endocytic tracer FM4-64 into *van3* cells (Naramoto et al., 2010). Together, these evidences suggest that the developmental function of VAN3 in vascular patterning and other processes in auxin-mediated development may be mediated, on a cellular level, through the regulation of CME at the PM.

Another ARF regulator, the ARF-GEF GNOM, which too has a broad function in auxin-mediated patterning processes, including embryonic patterning, vascular tissue patterning, and lateral root formation (Mayer et al., 1993; Steinmann, 1999; Koizumi et al., 2000; Geldner et al., 2003; Naramoto and Kyozuka, 2018; Verna et al., 2019), has also been localized to the PM, beside its site of action at the Golgi apparatus (Naramoto et al., 2010, 2014). Recent results indicate that the PM is the site of GNOM action relevant for its function in developmental patterning (Adamowski et al., 2021a). The localization of GNOM at the PM has been previously suggested to represent an activity in CME together with VAN3 (Naramoto et al., 2010), but the major form of GNOM localization to the PM are structures not containing clathrin, and *gnom* null mutant seedlings exhibit normal CME, arguing against this notion (Adamowski et al., 2021a). Taken together, there is currently no strong evidence that the cellular activity of GNOM, which translates into its function in developmental patterning, is associated with the regulation of CME.

These findings motivated us to address the functional connections of VAN3 to the clathrin-coated vesicle (CCV) formation machinery at the PM. In the present report, we re-evaluate the localization patterns of VAN3 at the PM by Total Internal Reflection Fluorescence (TIRF) microscopy, analyse the endocytic process in *van3* loss-of-function mutants by similar means, and conversely, test the requirement of clathrin for VAN3 function at the PM using novel inducible artificial microRNA lines silencing the clathrin coat component *CLATHRIN HEAVY CHAIN (CHC)*.

## Results

### VAN3 localizes to abundant and highly dynamic structures at the PM distinct from clathrin, DRP1A, or GNOM

To test the potential involvement of VAN3 in clathrin- and dynamin-mediated endocytosis, and a potential association with the GNOM-positive structures at the PM (Adamowski et al., 2021a), we studied the previously described double fluorescent protein marker lines of VAN3 with CLATHRIN LIGHT CHAIN (CLC), a component of the vesicular clathrin coat (*VAN3_pro_:VAN3-GFP 35S_pro_:CLC-mKO [mKusabira Orange])* and VAN3 with the ARF-GEF GNOM (*GNOM_pro_:GNOM-GFP VAN3_pro_:VAN3-mRFP*) (Naramoto et al., 2010), and additionally generated a new marker cross *VAN3_pro_:VAN3-GFP 35S_pro_:DRP1A-mRFP*, taking into account the physical interaction of VAN3 with the dynamin isoform DRP1A (Sawa et al., 2005). We captured TIRF time lapses of these double marker lines in the epidermis of etiolated hypocotyls and compared the localization patterns and dynamic behaviours of the fluorescent protein fusions. VAN3 was found to localize to the PM as discrete foci (Figure 1A, C, E), but they were clearly separate from sites of CCP formation marked by CLC or DRP1A (Figure 1A, C), and from the GNOM-positive PM structures (Figure 1E). Time lapse movies (Suppl. Movies 1-9) and kymographs representing the course of these time lapses (Figure 1B, D, F) demonstrated that VAN3 associated with the PM very briefly, appearing on kymographs as densely distributed spots, in contrast to CLC or DRP1A, which remained at the forming CCPs for relatively longer periods, and to GNOM, which often persisted in its specific structures for the whole length of the captured movies. The spots of VAN3 association with the PM in the kymographs were, again, not found to colocalize with traces of CLC, DRP1A, or GNOM. Taken together, these observations of VAN3 localization at the PM, using TIRF time lapses, show that VAN3 associates with the PM at sites different from CCPs, and different from those that recruit GNOM.

**Figure 1.**
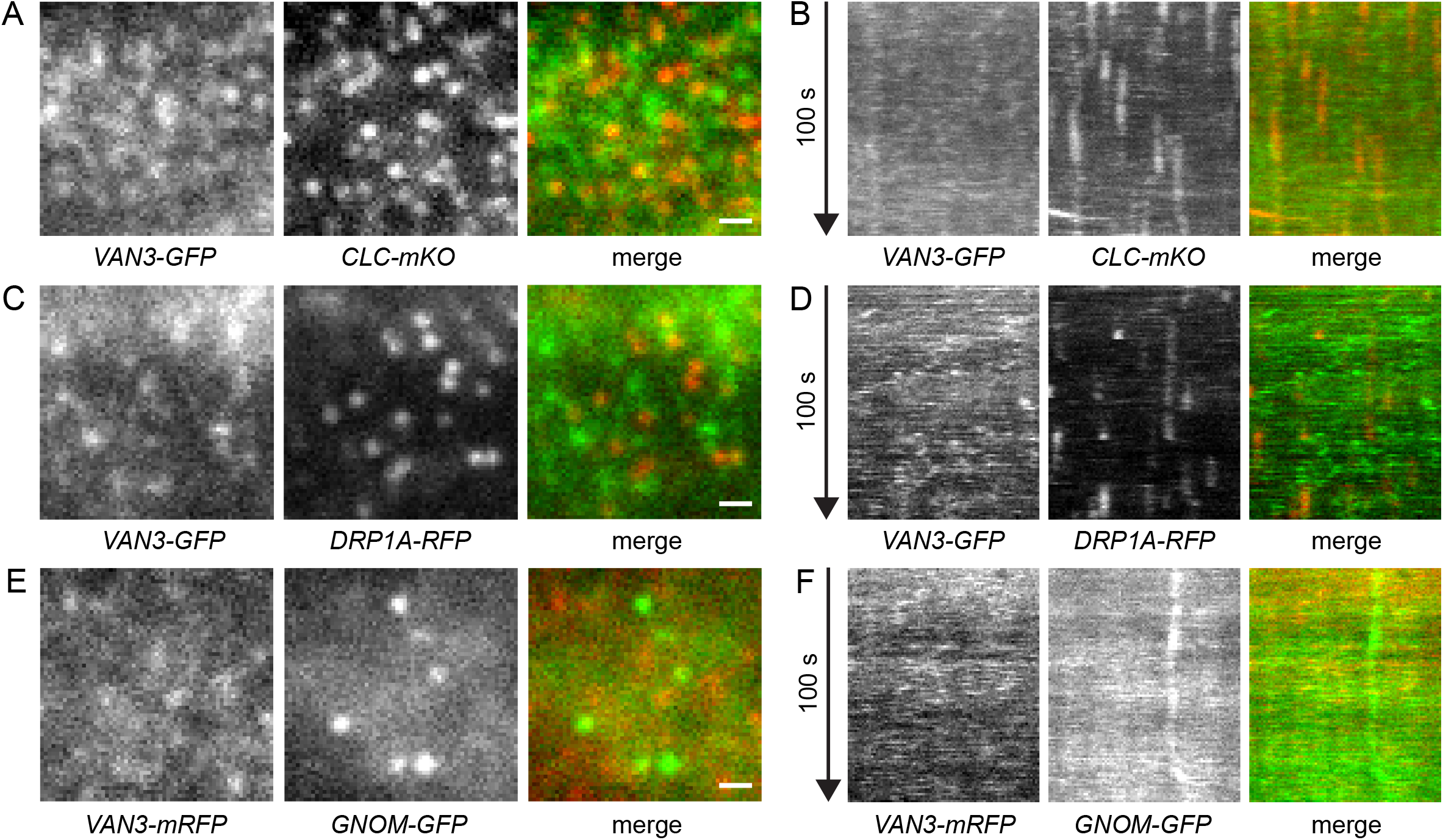
TIRF colocalization of VAN3 with CLC, DRP1A, and GNOM. Single frames (A, C, E) and kymographs (B, D, F) of TIRF movies capturing colocalization of VAN3-GFP with CLC-mKO (A, B), VAN3-GFP with DRP1A-RFP (C, D) and VAN3-mRFP with GNOM-GFP (E, F) at the PM of etiolated hypocotyl epidermis. VAN3 is present as discrete foci briefly associating with the PM at sites different from CLC, DRP1A, or GNOM. Scale bars – 1 μm.

### Clathrin-mediated endocytosis proceeds normally in *van3* knockout mutants

To explore further the potential functional connection of VAN3 to CME, we assessed the rates of CME in a *van3* loss of function mutant. We introduced fluorescent protein markers for the coat protein clathrin (*CLC2_pro_:CLC2-GFP;* Konopka et al., 2008) and for the TPLATE complex, presumably acting as a clathrin adaptor (*LAT52_pro_:TPLATE-GFP;* Gadeyne et al., 2014), into the *van3-1* knockout mutant background. We recorded TIRF time lapses of these CME markers in the epidermis of seedling roots of *van3-1* mutants and wild type controls to assess whether the endocytic process is defective as a result of loss of VAN3 function.

The density of CCPs marked by the presence of CLC2-GFP or TPLATE-GFP was quantified from a total sample of 15 to 17 roots of each genotype expressing each fluorescent marker. We found the density of CLC2-positive or TPLATE-positive foci to be the same in the wild type and in *van3-1* (Figure 2A, B, C, D), indicating a normal rate of endocytosis in the mutant. In time lapses, both CLC2-GFP and TPLATE-GFP occurred in two qualitatively distinguishable classes on a cell-to-cell basis (Figure 3 A, D; Suppl. Movies 10-15). Structures marked by CLC2-GFP or TPLATE-GFP were observed in some cells as predominantly dynamic entities (designated “short-type”), or, in other cells, as predominantly static structures that persisted throughout the 100s-long time lapses (designated “long-type”). Cells of a single root, visible in a single TIRF movie, typically all belonged to one type, but in similar experiments performed as part of other studies we found instances of neighbouring cells belonging to both distinct types (Suppl. Figure 1 and Suppl. Movie 16). We found no evident cause for the existence of these two apparent types of CCP dynamics: TIRF movies of all seedlings were captured in the same early elongation zone of the root, and we have no reasons to suspect tissue damage or stresses to be induced, especially in a large fraction of all imaged roots. In the sample captured for the present study, seedling roots with long-type CME events predominated in the wild type, while they were almost absent in the *van3-1* mutant (Figure 3B, E), but since the reason for the existence of these two distinct classes is unclear, it is also not clear whether their near absence in the mutant sample is meaningful. Putting aside this unexpected observation, when lengths of individual CME events were measured from kymographs of short-type cells of wild type and *van3-1*, the time courses of CCP formation were found to be distributed similarly (Figure 3 C, F), not indicative of any abnormalities in the formation of CCPs in the mutant.

**Figure 2.**
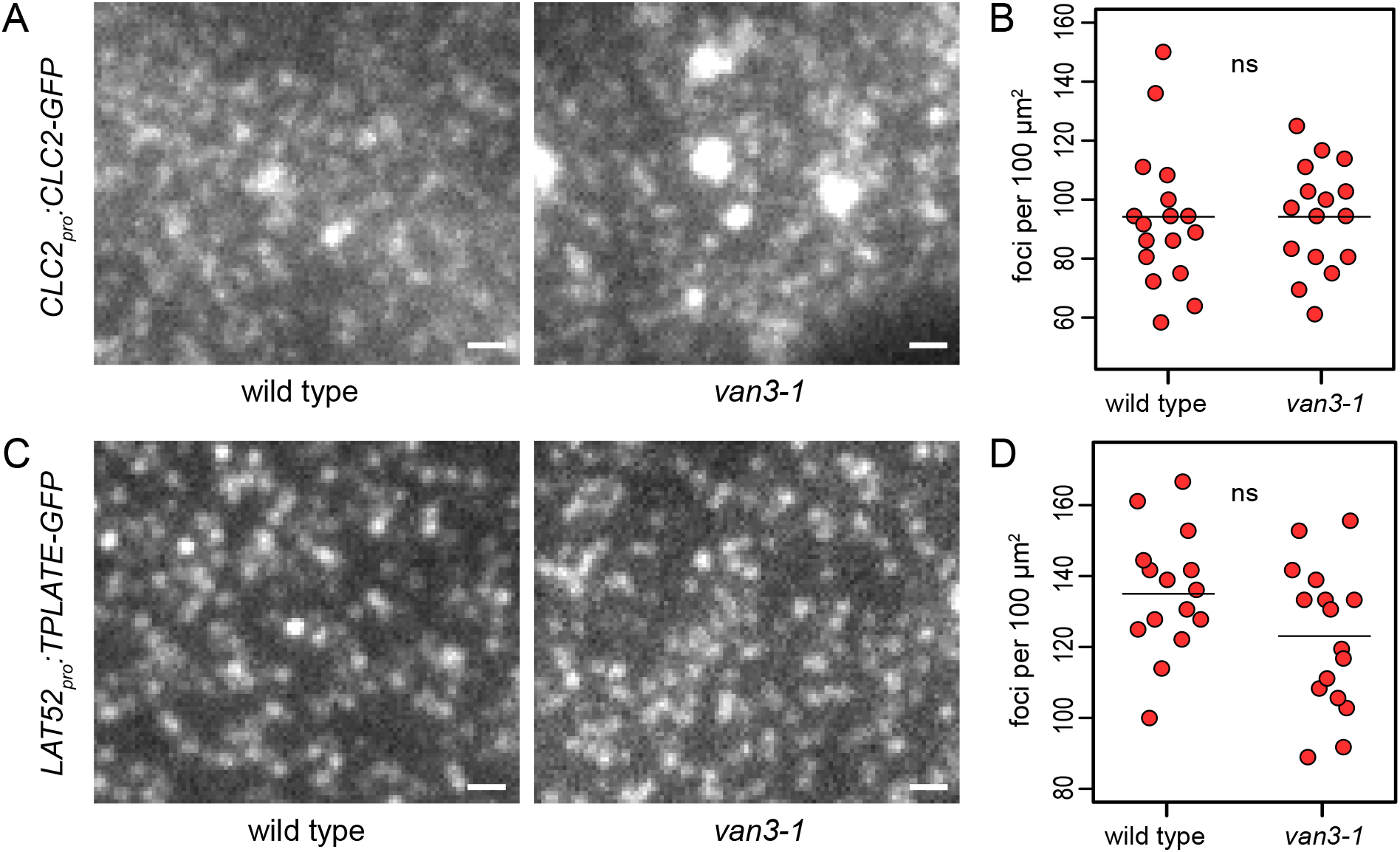
Density of endocytic events at the PM of *van3-1* mutants. Single frames of TIRF movies capturing the dynamics of CLC2-GFP (A) and TPLATE-GFP (C) at the PMs of seedling root epidermis in *van3-1* and wild type controls. Densely distributed spots represent single CCPs forming at the PM surface. The large structures in (A) are clathrin-positive TGN/EEs in close proximity to the PM. Scale bars – 1 μm. Strip charts (B, D) show quantifications of endocytic event densities at the PM, each data point representing a measurement from a single root. Horizontal bars represent data means. CLC2-GFP wild type: 94 ± 28 foci per 100 μm^2^, n=17; *van3-1*: 94 ± 18 foci per 100 μm^2^, n=16. TPLATE-GFP wild type: 135 ± 17 foci per 100 μm^2^, n=15; *van3-1*: 123 ± 20 foci per 100 μm^2^, n=17. Values were compared using *t* tests, ns – not significant.

**Figure 3.**
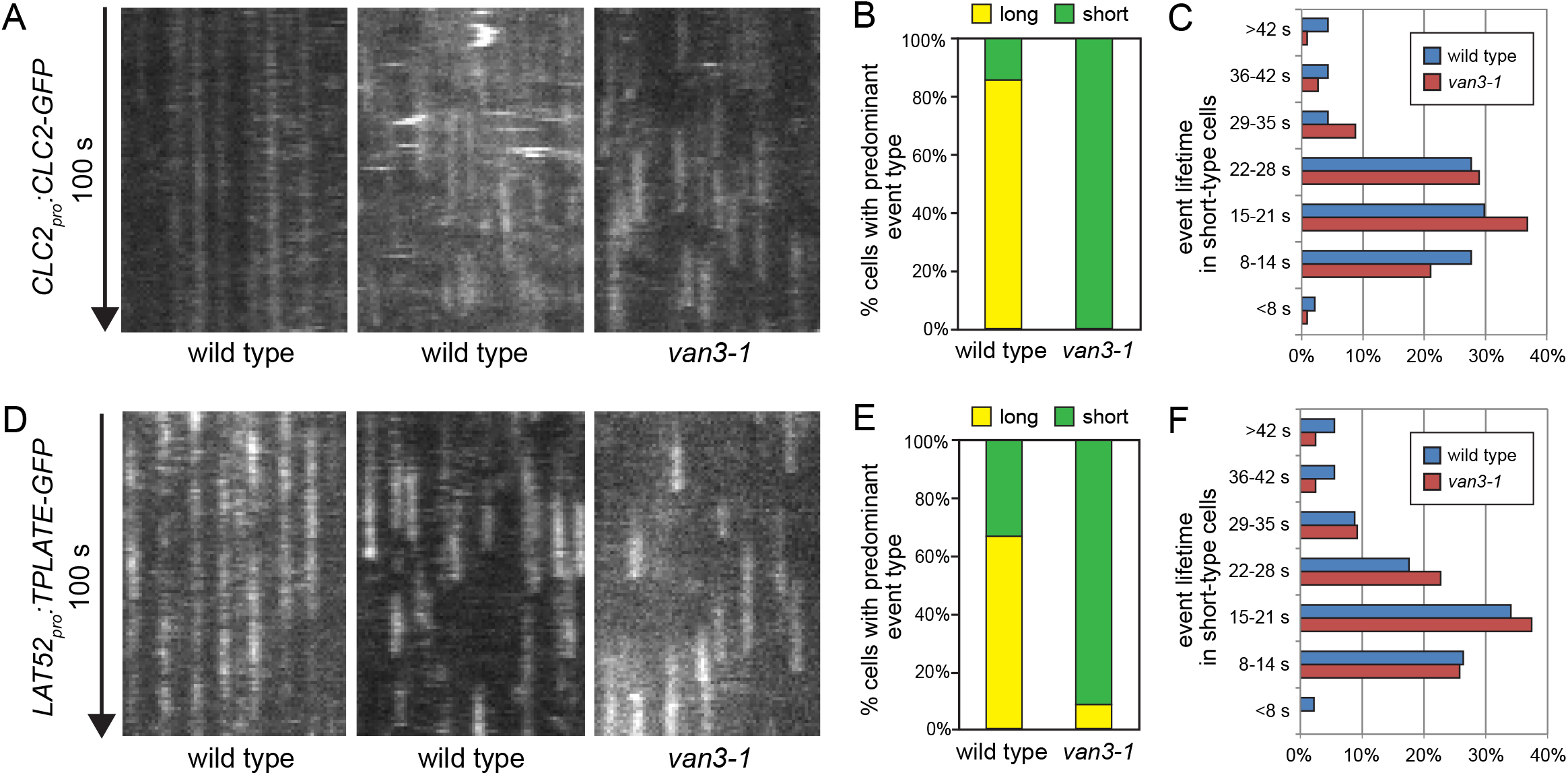
Temporal dynamics of endocytic events at the PM of *van3-1* mutants. Kymographs of TIRF movies capturing the dynamics of CLC2-GFP (A) and TPLATE-GFP (D) at the PMs of seedling root epidermis in *van3-1* and wild type controls. Images of wild type show cells with predominantly long-type endocytic events (left panel) and with predominantly short-type endocytic events (middle panel). Graphs (B, E) show sum frequencies of cells with predominantly long-type and short-type endocytic events in all captured TIRF movies. Histograms (C, F) show the distribution of lifetimes of single endocytic events in short-type cells of wild type and *van3-1* mutants. CLC2-GFP wild type n=47, *van3-1* n=114; TPLATE-GFP wild type n=91, *van3-1* n=163.

Taken together, TIRF imaging of CCP marker proteins in *van3-1* indicates that CME proceeds normally in terms of the density of initiated vesicle formation events, and that the time course of CCP development is normal in cells with short-type dynamics. The unexpected finding that CME can proceed with slow dynamics, and the near absence of this phenomenon in *van3-1*, are at present unexplained. Together with the lack of co-localization of VAN3 with CLC or DRP1A at the PM, we find no compelling evidence that VAN3 participates in, or is required for, CME.

### Clathrin is required for VAN3 localization at the PM, likely due to clathrin’s function in endocytosis but not exocytosis

The data presented above argue against an involvement of VAN3 in endocytosis. To further explore potential links between VAN3 and clathrin, we asked the converse question: does VAN3 require clathrin for its normal localization? We employed the recently developed transgenic lines of *A. thaliana* expressing estradiol-inducible artificial microRNA lines silencing *CHC*. After 2 days of transgene induction, silencing of *CHC* leads to an inhibition of CME, and causes defects in exocytosis, which are not associated with a general disruption of the TGN compartment, as exocytic cargoes still traffic through this compartment, and are re-routed to the vacuole (Adamowski et al., 2021b). Using *XVE»amiCHCa* introduced into *GNOM_pro_:GNOM-GFP VAN3_pro_:VAN3-mRFP* (Adamowski et al., 2021a), we now analysed the effects of clathrin silencing on VAN3 localization. We employed CLSM (Confocal Laser Scanning Microscopy) in the epidermis of seedling root apical meristems (RAMs), approximately 48 h after induction of amiRNA expression by β-estradiol. A moderate reduction in the overall signal levels of VAN3-mRFP, indicative of an effect on protein expression, was observed (Suppl Fig. 2A). With regard to VAN3 localization at the PM, we observed a clear loss of signal association with the PM, and VAN3 now localized predominantly in the cytosol (Figure 4A). This suggests that clathrin function is required for normal association of VAN3 to the PM, and as such, presumably for its normal activity, possibly in an indirect manner.

**Figure 4.**
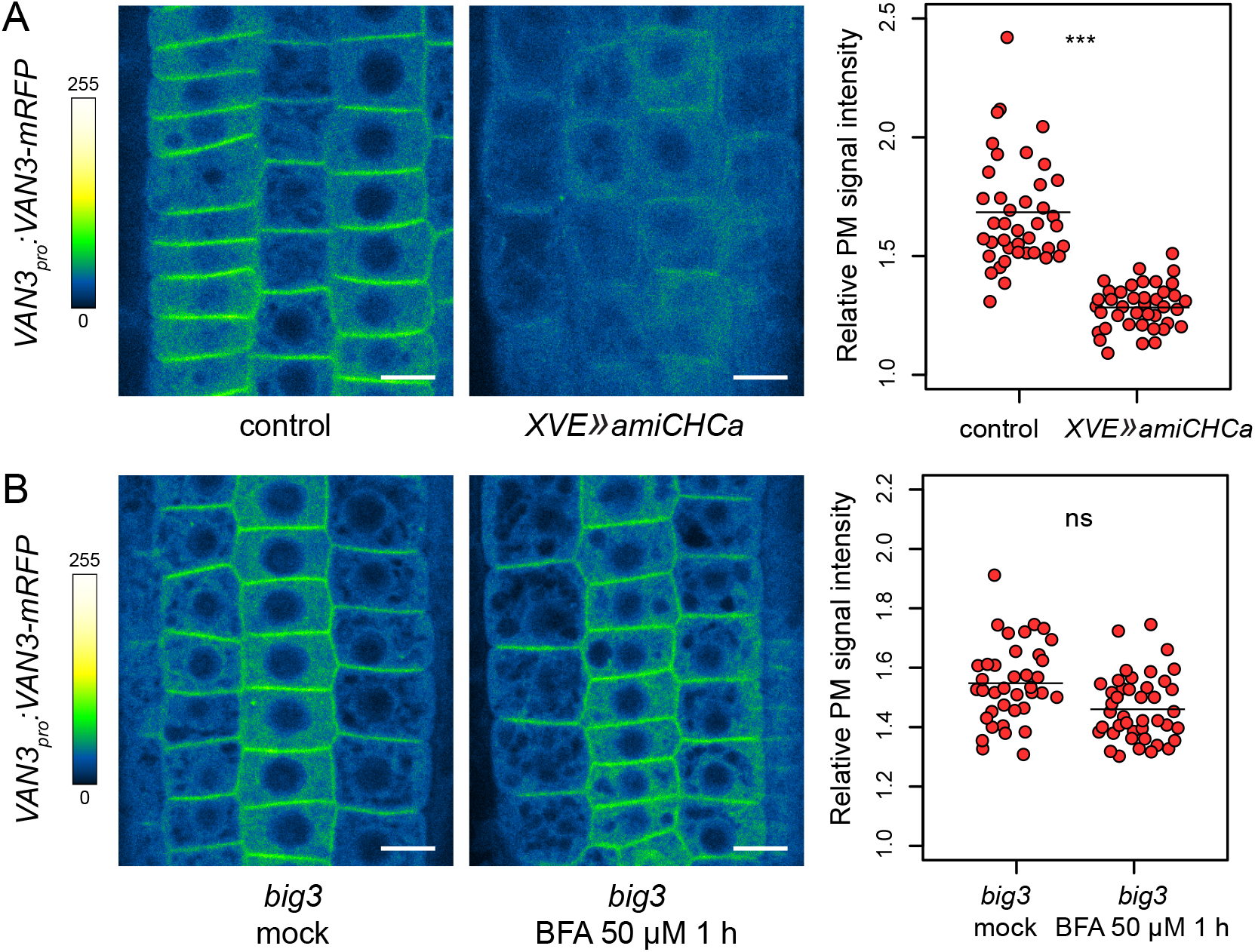
Dependence of VAN3 targeting to the PM on clathrin-mediated endocytosis. CLSM images of VAN3-mRFP in seedling RAMs following inhibition of clathrin function by post-translational silencing of *CHC* for approximately 48 h (A) and following inhibition of exocytosis from the TGN by the application of BFA at 50 μM for 1 h in *big3* mutant background (B). Scale bars – 10 μm. Strip charts show intensities of VAN3-mRFP signals at the PMs relative to total root meristem signals, each data point represents one root. Horizontal bars represent data means. (A) Control 1.68 ± 0.23, n=41, *XVE»amiCHCa* 1.28 ± 0.09, n=40. (B) *big3* mock 1.55 ± 0.13, n=38, *big3* BFA 50 μM 1 h 1.46 ± 0.11, n=42. Values were compared using *t* tests, *** P<0.0001; ns – not significant.

Silencing of clathrin with amiRNA affects clathrin function unselectively, down-regulating CME and clathrin-dependent exocytosis both (Adamowski et al., 2021b). We conceived an experiment aimed at distinguishing whether it is CME, or clathrin-dependent exocytosis, that is required for VAN3 association with the PM. A synthesis of previously published data indicates a scenario where clathrin-dependent exocytosis may occur through vesicles coated with the clathrin adaptor ADAPTOR PROTEIN-1 (AP-1), whose formation is dependent on ARF small GTPases activated by BIG class ARF-GEFs at the TGN. Defects in secretion are observed in *ap-1* loss of function (Park et al., 2013) and following BIG ARF-GEF inhibition (Richter et al., 2014). In non-plant systems, the clathrin adaptor AP-1 is known to be an ARF effector, i.e. a protein recruited to membranes by activated ARFs (Paczkowski et al., 2015), and this is likely the case in plants as well, since AP-1 loses binding to TGN membranes, and relocates to the cytosol, when BIG ARF-GEFs are inhibited (Richter et al., 2014). Following this reasoning, clathrin-dependent exocytosis at the TGN can be blocked selectively, without affecting CME, when BIG class ARF-GEFs are inhibited, as this will prevent ARF activation leading to AP-1 recruitment and formation of clathrin-coated pits at the TGN. BIGs can be inhibited effectively by the combination of *big3* mutation, and of chemical inhibition of remaining homologues with Brefeldin A (BFA), due to the fact that BIG3 contains a SEC7 domain variant that is resistant to BFA (Richter et al., 2014). Thus, to test if TGN-localized clathrin is required for VAN3 localization at the PM, we employed a *GNOM_pro_:GNOM-GFP VAN3_pro_:VAN3-mRFP* line crossed into *big3* mutant background (Adamowski et al., 2021a) and applied BFA. The result of this experiment is negative: Following a treatment with BFA at 50 μM for 1h in *big3* mutant background, VAN3 is normally retained at the PM (Figure 4B). Expectedly, the result is the same in wild type background, where effects of BFA are limited due to the activity of BIG3 (Suppl. Figure 2B). The experiment argues against a requirement of BIG-mediated exocytosis, which, as explained, is likely identical to clathrin-dependent exocytosis, for VAN3 localization at the PM. By exclusion, this indicates that the loss of VAN3 from the PMs observed when clathrin function is down-regulated unselectively in *XVE»amiCHCa*, is an outcome of a defective CME.

## Discussion

The developmentally important ARF-GAP VAN3 has been proposed to act from the PM in a process involving the regulation of CME (Naramoto et al., 2010; Naramoto and Kyozuka, 2018). In the present study, using now available state-of-the-art TIRF imaging we scrutinize the character of VAN3 ARF-GAP presence at the PM. Functional fluorescent protein fusions of VAN3, whose localization at the PM has been asserted as the relevant site of VAN3 action (Naramoto and Kyozuka, 2018), associate with the PM transiently and at sites distinct from the endocytic machinery. The higher resolution imaging presented here does not support the previous observations identifying partial co-localization between VAN3 and clathrin (Naramoto et al. 2010) likely due to the use of time lapses, and kymographs, which both help identify signals representing meaningful structures containing the fluorescent reporters. The previously described physical interaction between VAN3 and the dynamin isoform DRP1A (Sawa et al., 2005) also does not translate to *in vivo* associations at the PM, and as such, the relevance of this interaction in VAN3 activity remains unclear. Given the lack of recruitment to CCPs, it is also an open question in which sense the potential curvature-sensing properties of the BAR domain of VAN3 may contribute to its function at the PM.

Consistently with VAN3 being present at structures distinct from CCPs, we find that CME proceeds normally in a *van3* knockout mutant, both in terms of densities of initiated endocytic events, and time courses of their development, when cells with short-type dynamics are considered. The identification of long-type endocytic dynamics, observed with CLC2-GFP and TPLATE-GFP markers, remains a challenging observation, as there is no clear reason why CCPs should be formed with such slow dynamics, or indeed, be apparently arrested, in otherwise normally functioning cells. A similar phenomenon was previously observed with a dynamin marker DRP1C-GFP, but in conditions of CME inhibition caused by the overexpression of a putative CCV uncoating factor AUXILIN-LIKE1 (Adamowski et al., 2018). The observation of long-type dynamics in wild type cells should be further verified by functional fluorescent protein markers expressed in complemented mutant backgrounds. While this phenomenon was almost never observed in the *van3* mutant, we refrain at present from associating it with VAN3 function due to the general difficulty in understanding the nature of this unexpected observation.

While the localization of VAN3 at the PM, as well as measurements of CME in *van3*, point to VAN3 activity different than the regulation of CME, we identified a converse dependence, where VAN3 localization at the PM requires clathrin function. The comparison of *XVE»amiCHCa* with *big3* mutants treated with BFA, where AP-1-related clathrin function at the TGN is inhibited selectively, indicates that the function of clathrin at the PM specifically is most likely required to maintain VAN3 presence. This dependency may be indirect: Possibly, in analogy to the control of VAN3 activity by VAB/FKD1, CVP2, and CVL1, regulators which influence the binding of VAN3 to membrane phospholipids through its PH domain (Naramoto et al., 2009; Carland and Nelson, 2009), an influence of functional CME on lipid composition of the PM as a whole, may be responsible.

While our study narrows down future investigations by the exclusion of VAN3’s involvement in CME, the nature of VAN3-containing foci at the PM, and the activity of VAN3 at the PM through which its developmental function is carried out, remains unidentified. Similar challenges arise from the recent studies of the ARF-GEF GNOM. Just like VAN3, GNOM is an ARF regulator acting from the PM, required for developmental patterning, and whose function is not associated with the control of CME. GNOM acts from relatively sparse, large, and usually stable structures at the PM, different from those where VAN3 localizes, but whose nature is equally unknown (Adamowski et al., 2021a). The understanding of the mode of action of ARF small GTPase machinery in developmental patterning presents yet another challenge: only a single member of the ARF family in *A. thaliana*, ARFB1a, localizes to the PM (Matheson et al., 2008; Singh et al., 2018; Adamowski et al., 2021c), but even high order loss of function mutants, including *arfb1a* and several family members most similar to it, do not exhibit any visible phenotypes, let alone developmental patterning defects reminiscent of *van3* or *gnom* (Adamowski et al., 2021c; Singh et al., 2018). Thus, the molecular mode of action of the ARF regulators involved in developmental patterning is at present far from being elucidated.

## Materials and methods

### Plant material

The following previously described *A. thaliana* lines were used in this study: *van3-1* (Koizumi et al., 2000)*, CLC2_pro_:CLC2-GFP* (Konopka et al., 2008), *LAT52_pro_:TPLATE-GFP RPS5A_pro_:AP2A1-TagRFP tplate* (Gadeyne et al., 2014), *VAN3_pro_:VAN3-GFP, VAN3_pro_:VAN3-GFP 35S_pro_:CLC-mKO, GNOM_pro_:GNOM-GFP VAN3_pro_:VAN3-mRFP* (Naramoto et al., 2010), *35S_pro_:DRP1A-mRFP* (Mravec et al., 2011), *XVE»amiCHCa GNOM_pro_:GNOM-GFP VAN3_pro_:VAN3-mRFP, big3 GNOM_pro_:GNOM-GFP VAN3_pro_:VAN3-mRFP* (Adamowski et al., 2021a). Lines generated as part of this study: *van3-1 CLC2_pro_:CLC2-GFP, van3-1 LAT52_pro_:TPLATE-GFP RPS5A_pro_:AP2A1-TagRFP, VAN3_pro_:VAN3-GFP 35S_pro_:DRP1A-mRFP*. *van3-1* mutants were genotyped by sequencing of PCR products amplified with primers van3-1-F: TGCAAACAATAAGGCAAGGTT and van3-1-R: TGATGCAATAACCCCAGTGA.

### *In vitro* cultures of *Arabidopsis* seedlings

Seedlings were grown in *in vitro* cultures on half-strength Murashige and Skoog (½MS) medium of pH=5.9 supplemented with 1% (w/v) sucrose and 0.8% (w/v) phytoagar at 21°C in 16h light/8h dark cycles with Philips GreenPower LED as light source, using deep red (660nm)/far red (720nm)/blue (455nm) combination, with a photon density of about 140μmol/(m^2^s) +/- 20%. Petri dishes for TIRF imaging in hypocotyls of etiolated seedlings were initially exposed to light for several hours and then wrapped in aluminium foil until the time of imaging.

### Chemical treatments

BFA (Sigma-Aldrich) was solubilized in DMSO to 50 mM stock concentration and added to liquid ½MS media for treatments. Beta-estradiol (Sigma-Aldrich) was solubilized in 100% ethanol to 5 mg/mL stock concentration and added to ½MS media during preparation of solid media to a final concentration of 2.5 μg/mL. Induction of *XVE»amiCHCa* was performed approximately 48 h before CLSM imaging by transferring 3 day old seedlings to media supplemented with beta-estradiol.

### Total Internal Reflection Fluorescence microscopy

A sub-apical region of excised hypocotyls of 3d old dark-grown seedlings was used for fluorescent marker colocalization. Early elongation zone of root meristems in excised ~1 cm long root tip fragments from 7d old light-grown seedlings was used for imaging of CME markers in *van3-1* mutants. Phenotypically wild type plants from segregating *van3-1* populations, distinguished by normal growth rates, were used as controls. Imaging was performed with Olympus IX83 TIRF microscope, using a 100X TIRF lens with an additional 1.6X magnification lens in the optical path. Time lapses of 100 frames at 1 s intervals, with exposure times of 200 ms, were taken. In colocalization studies, green and red channels were captured sequentially.

### Confocal Laser Scanning Microscopy

5 d old seedlings were used for live imaging with Zeiss LSM800 confocal laser scanning microscope with 20X lens.

### Analysis of live imaging data

Fiji (https://imagej.net/Fiji) was used for all quantifications. CLC2-GFP and TPLATE-GFP foci were counted in square regions of 36 μm^2^ taken from a single still frame of each captured TIRF movie. Lifetimes of CLC2-GFP and TPLATE-GFP were measured in kymographs extracted from each captured TIRF movie where cells with short-type dynamics were found.

VAN3-mRFP PM signal intensities were measured as mean grey value of a line of 5 pixel width drawn over multiple PMs in each CLSM image. Total signal intensity was measured as mean grey value of a rectangle covering whole RAM surface visible in an image. Ratios of the two measurements were counted and compared between conditions.

## Accession numbers

Sequence data from this article can be found in the GenBank/EMBL libraries under the following accession numbers: AGD3/VAN3/FKD2/SFC (AT5G13300), DRP1A (AT5G42080), GNOM (AT1G13980), BIG3 (AT1G01960).

## Supplemental Material

Suppl. Figure 1. Adjacent short-type and long-type cells expressing TPLATE-GFP

Suppl. Figure 2. Additional data on VAN3 localization

Suppl. Movie 1. TIRF colocalization of VAN3-GFP and CLC-mKO, green channel

Suppl. Movie 2. TIRF colocalization of VAN3-GFP and CLC-mKO, red channel

Suppl. Movie 3. TIRF colocalization of VAN3-GFP and CLC-mKO, merged channels

Suppl. Movie 4. TIRF colocalization of VAN3-GFP and DRP1A-RFP, green channel

Suppl. Movie 5. TIRF colocalization of VAN3-GFP and DRP1A-RFP, red channel

Suppl. Movie 6. TIRF colocalization of VAN3-GFP and DRP1A-RFP, merged channels

Suppl. Movie 7. TIRF colocalization of VAN3-mRFP and GNOM-GFP, red channel

Suppl. Movie 8. TIRF colocalization of VAN3-mRFP and GNOM-GFP, green channel

Suppl. Movie 9. TIRF colocalization of VAN3-mRFP and GNOM-GFP, merged channels

Suppl. Movie 10. TIRF time lapse of long-type CLC2-GFP in the wild type

Suppl. Movie 11. TIRF time lapse of short-type CLC2-GFP in the wild type

Suppl. Movie 12. TIRF time lapse of short-type CLC2-GFP in *van3-1*

Suppl. Movie 13. TIRF time lapse of long-type TPLATE-GFP in the wild type

Suppl. Movie 14. TIRF time lapse of short-type TPLATE-GFP in the wild type

Suppl. Movie 15. TIRF time lapse of short-type TPLATE-GFP in *van3-1*

Suppl. Movie 16. TIRF time lapse of TPLATE-GFP in the wild type with adjacent short-and long-type cells

## Acknowledgements

This work was supported by the Austrian Science Fund (FWF): I 3630-B25.

---

**Supplementary Figure 1.**
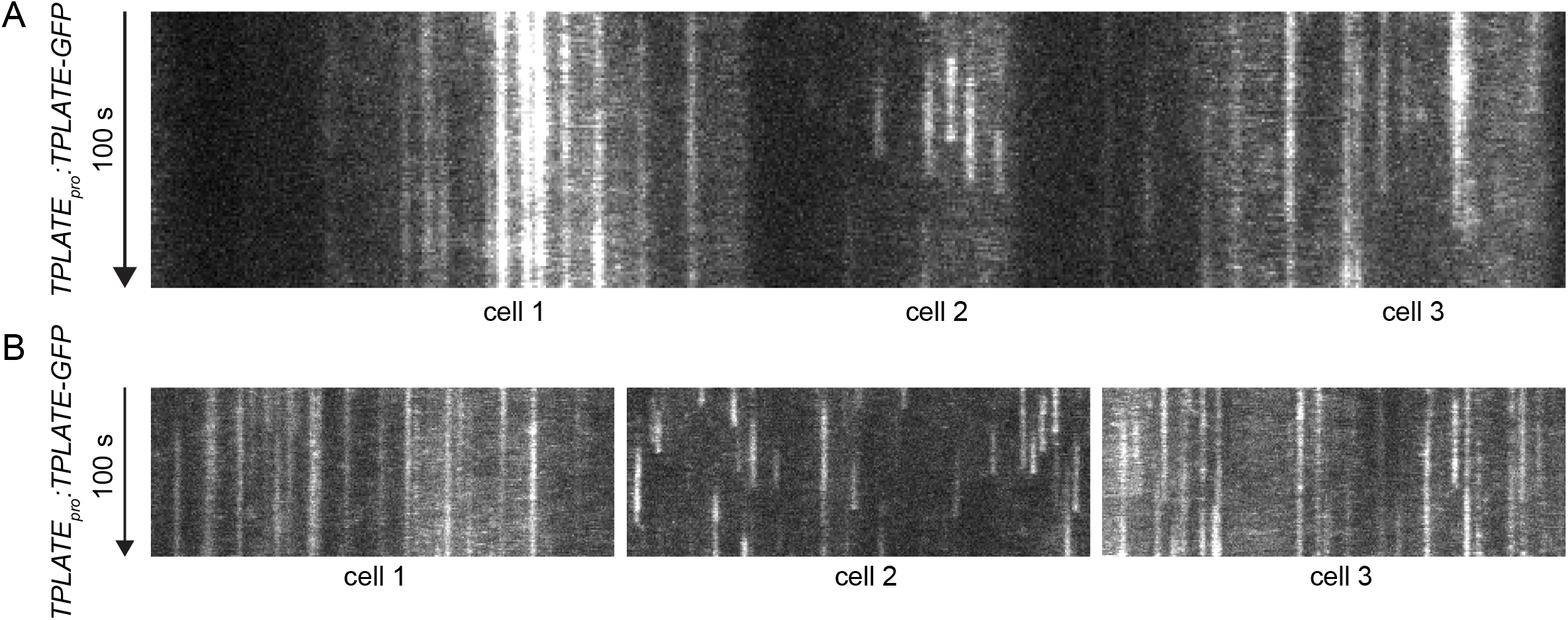
Adjacent short-type and long-type cells expressing TPLATE-GFP. Kymographs of a single TIRF movie (Suppl. Movie 16) showing cells with long-type TPLATE-GFP dynamics (cell 1 and 3) surrounding a cell with short-type TPLATE-GFP dynamics (cell 2), captured in the early elongation zone of a seedling root from a wild type plant as part of another study. (A) A single horizontal line section through the movie, (B) Additional vertical line sections through each cell.

**Supplementary Figure 2.**
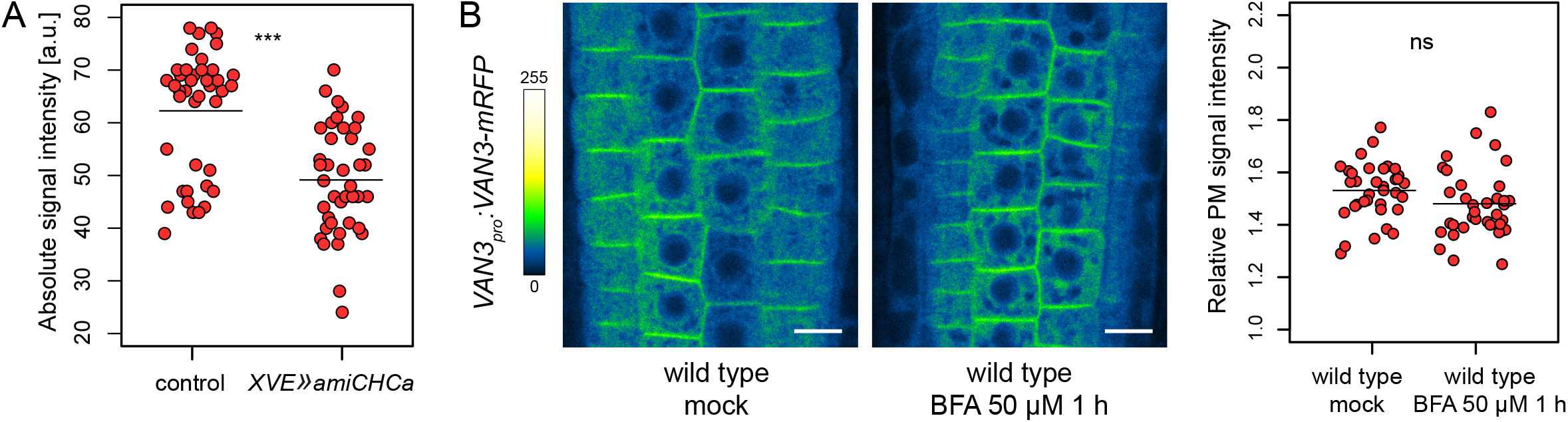
Additional data on VAN3 localization. (A) Absolute signal intensities of VAN3-mRFP in seedling RAMs following inhibition of clathrin function by post-translational silencing of *CHC* for approximately 48 h and in control conditions. Each data point represents one root. Horizontal bars represent data means. Control 62.24 ± 11.47, n=41, *XVE»amiCHCa* 49.13 ± 10.22, n=40. Values were compared using a *t* test, *** P<0.0001 (B) Control CLSM images of VAN3-mRFP in seedling RAMs following BFA treatment in the wild type. Scale bars – 10 μm. Strip chart shows intensities of VAN3-mRFP signals at the PMs relative to total root meristem signals, each data point represents one root. Horizontal bars represent data means. mock 1.53 ± 0.11, n=35, BFA 50 μM 1 h 1.48 ± 0.13, n=37. Values were compared using a *t* test, ns – not significant.

